# Beyond resource selection: emergent spatio-temporal distributions from animal movements and stigmergent interactions

**DOI:** 10.1101/2022.02.28.482253

**Authors:** Jonathan R. Potts, Valeria Giunta, Mark A. Lewis

## Abstract

A principal concern of ecological research is to unveil the causes behind observed spatio-temporal distributions of species. A key tactic is to correlate observed locations with environmental features, in the form of resource selection functions or other correlative species distribution models. In reality, however, the distribution of any population both affects and is affected by those surrounding it, creating a complex network of feedbacks causing emergent spatio-temporal features that may not correlate with any particular aspect of the underlying environment. Here, we study the way in which the movements of populations in response to one another can affect the spatio-temporal distributions of ecosystems. We construct a stochastic individual-based modelling (IBM) framework, based on stigmergent interactions (i.e. organisms leave marks which cause others to alter their movements) between and within populations. We show how to gain insight into this IBM via mathematical analysis of a partial differential equation (PDE) system given by a continuum limit. We show how the combination of stochastic simulations of the IBM and mathematical analysis of PDEs can be used to categorise emergent patterns into homogeneous vs. heterogeneous, stationary vs. perpetually-fluctuating, and aggregation vs. segregation. In doing so, we develop techniques for understanding spatial bifurcations in stochastic IBMs, grounded in mathematical analysis. Finally, we demonstrate through a simple example how the interplay between environmental features and between-population stigmergent interactions can give rise to predicted spatial distributions that are quite different to those predicted purely by accounting for environmental covariates.

## 1 Introduction

Understanding the processes behind the spatial distributions of animal populations has been a core concern of ecological research throughout its history (Elton, 2001; Nathan *et al*., 2008). Today, the need to manage the effects of rapid anthropogenic actions on ecosystems makes predictive tools for spatial ecology more important than ever (Azaele *et al*., 2015; Maris *et al*., 2018). However, spatial ecology is complicated by the fact that the distribution of a population of organisms will affect the distributions of those populations that surround it, and also be affected by these populations (Morales *et al*., 2010; Ovaskainen & Abrego, 2020). This generates a complex network of feedbacks between the constituent populations of an ecosystem, causing spatio-temporal patterns that can be difficult to predict, and impossible without the correct mathematical and computational tools linking process to pattern (May, 2019; Potts & Lewis, 2019).

There are two principal processes by which space use can be affected by interactions between populations (we use the word ‘population’ loosely, referring to anything ranging from a small group such as a territorial unit or herd through to an entire species). First, interactions can affect *demographics*, i.e. birth- and death-rates. This can be, for example, through predator-prey interactions or competition for resources, both of which are well-known to have non-trivial effects on both the overall demographic dynamics and the spatial distribution of species (Holmes *et al*., 1994; Tilman *et al*., 1997; Okubo & Levin, 2001; Cantrell & Cosner, 2004; Lewis *et al*., 2013, 2016).

Second, for mobile organisms, population interactions can affect the *movement* of individuals (Mitchell & Lima, 2002; Vanak *et al*., 2013; Breed *et al*., 2017; Matthews *et al*., 2020). It is well-known, from the mathematical literature, that the two processes of demographics and movement can combine to affect spatial distribution patterns in non-trivial ways, as exemplified by studies of cross-diffusion and prey-taxis (Shigesada *et al*., 1979; Lee *et al*., 2009; Gambino *et al*., 2013; Potts & Petrovskii, 2017; Han & Dai, 2019; Haskell & Bell, 2020). These studies typically model movement and demographics in the same system of equations (usually partial differential equations), implying that the movements are occurring on the same spatiotemporal scale as the demographics. Therefore the movements considered in such studies are usually dispersal events. However, many animal populations can make significant movements to rearrange themselves in space over timescales where births and deaths are negligible (Moorcroft *et al*., 2006; Vanak *et al*., 2013; Ellison *et al*., 2020). This particularly applies to larger animals, such as birds, mammals, and reptiles, who have great capability for movement but may only reproduce at a particular time of the year (e.g. spring). Therefore it is important to understand how movement processes alone may affect spatio-temporal population patterns (Potts & Lewis, 2019).

Spurred by rapid improvements in animal tagging technology, the empirical study of movement has surged, with data being gathered at ever higher resolutions (Williams *et al*., 2020). Furthermore, an increasing number of studies are measuring animal interactions via the co-tagging of multiple animals and new techniques for decoding the resulting information (Vanak *et al*., 2013; Potts *et al*., 2014c; Schlägel *et al*., 2019). A key goal of movement ecology is to understand animal space use, so the question of how fine-grained movement and interaction processes upscale to broader spatio-temporal patterns is gaining significant methodological attention (Avgar *et al*., 2016; Signer *et al*., 2017; Potts & Schlägel, 2020). However, to make predictions requires a theoretical understanding of how movements mediated by between-population interactions affect space use. Our principal aim here is to provide the theoretical framework for answering such questions.

To this end, we construct a general and extensible individual-based model (IBM) of movements and interactions between multiple populations. We assume that animals, left alone on the landscape, will have some sort of movement process allowing them to embark on daily activities such as foraging. We model this very simply as a nearest-neighbour lattice random walk (Okubo & Levin, 2001; Codling *et al*., 2008). This is a foundational movement model, which can be readily extended if one is interested in the finer details of foraging activity.

In this study, however, our focus is on the interactions between individuals and populations. For this, we assume that, as individuals move, they leave a trace of where they have been on the landscape, which could be in the form of scent, visual or olfactory marks, feces or a simply a trail. These marks decay over time if the area is not revisited. Consequently, each population leaves a distribution of such marks on the landscape, which changes over time as the constituent individuals move about. Individuals of a population alter their movement according to the presence or otherwise of marks, both from their own population and from others.

This process of leaving marks that cause others to alter their movement is called stigmergy, and has been studied in various contexts, including collective animal movement and territorial formation (Theraulaz & Bonabeau, 1999; Giuggioli *et al*., 2013; White *et al*., 2020). For any given pair of populations, *A* and *B*, one could either have mutual avoidance (where individuals from *A* avoid the marks of *B* and *B* avoid those of *A*), mutual attraction (individuals from *A* and *B* are attracted to the marks of one another), or pursuit-and-avoidance (individuals from *A* are attracted to marks of *B* but those from *B* avoid the marks of *A*). These combine into a network of stigmergent interactions that together determine the overall spatio-temporal distribution of the constituent populations (Figure 1). Our model is a generalisation of previous models of territory formation from stigmergent interactions (Giuggioli *et al*., 2011, 2013; Potts *et al*., 2012). However, these previous models were restricted to mutual avoidance processes and typically had only one individual per ‘population’ (recall, we are using ‘population’ quite generically here and could mean anything from a territorial unit to a larger group to a whole species, depending on context).

**Fig. 1.**
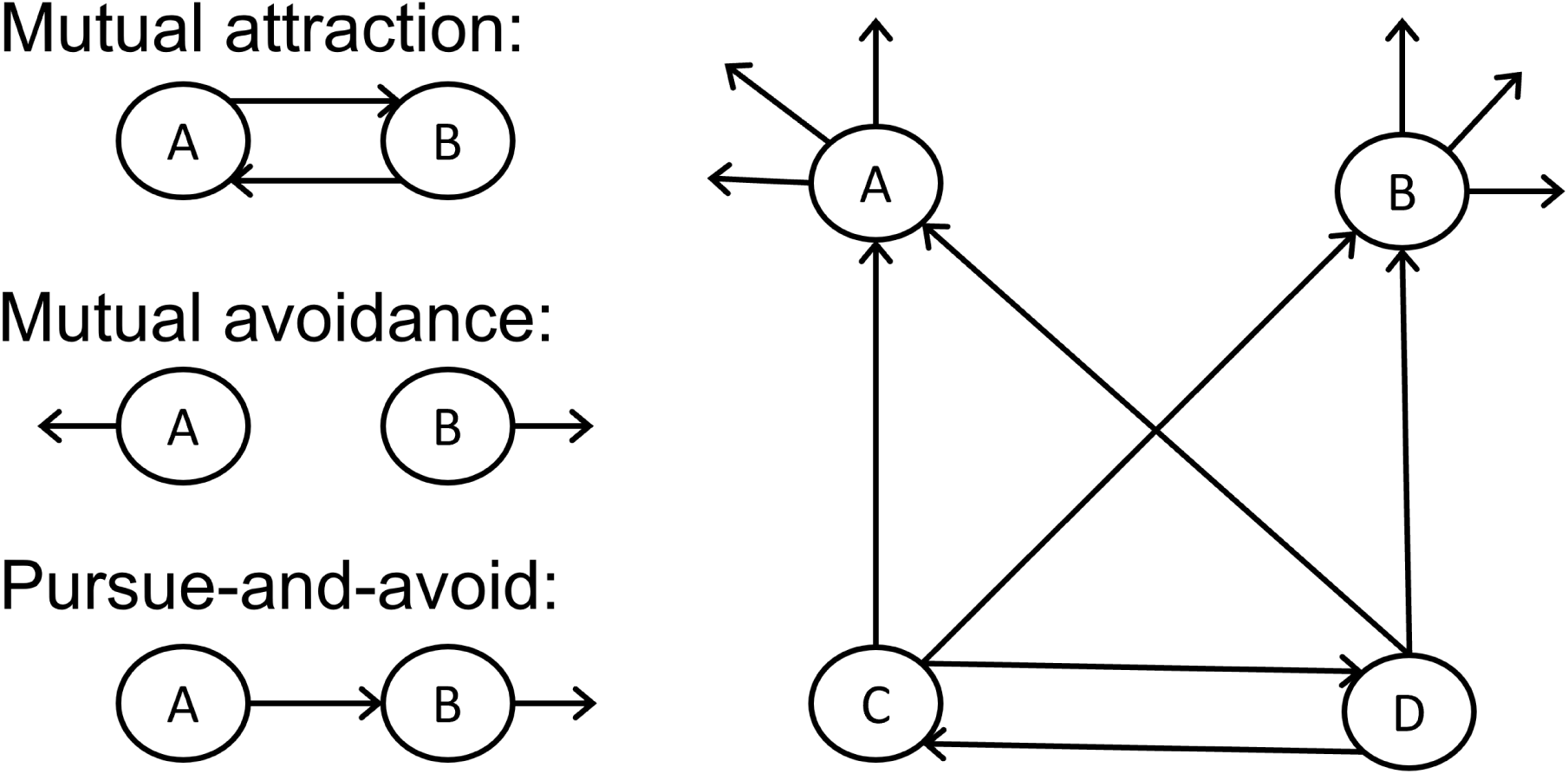
Schematic diagram of stigmergent interactions. The left-hand side shows the three possible pairwise interactions between two populations. On the right is an example network built from these interactions. One might imagine that A and B are competing prey being predated by mutualistic predators C and D.

As well as stochastic simulation analysis, we also examine the continuum limit of our IBM model in space and time (i.e. as the lattice spacing and time step go to zero). We construct the IBM so that this limit is a system of partial differential equations (PDEs) studied previously in Potts & Lewis (2019). This formal connection between IBM and PDE enables us to use the mathematical tools of PDE analysis to gain insight into the expected behaviour of the IBM, which we can verify through simulation. The resulting techniques allow us to use PDE analysis as a starting-point for exploring IBM models. This is valuable because PDEs are amenable to mathematical analysis, enjoying a huge history of analytic techniques (Evans, 2010; Murray, 2012). However, IBMs are closer to reality and may be more amenable to extensions that incorporate further realism beyond what is studied here (for example, realistic movement processes based on life history needs such as foraging and tending to young). Such formal connections between IBMs and PDEs are powerful as they enable the best of both worlds: combining rigorous mathematical analysis with realistic modelling.

Finally, we explain how to account for landscape heterogeneity in our model, through coupling our IBM to a step selection function (Fortin *et al*., 2005; Potts *et al*., 2014a; Avgar *et al*., 2016). We illustrate this with a simple example of two co-existing populations competing for the same resource, inspired by wolf-coyote coexistence in the Greater Yellowstone Ecosystem (Arjo & Pletscher, 2000). We investigate how the inclusion of interactions between and within the populations combine with the heterogeneous landscape. We show how this combination can cause emergent spatio-temporal patterns that cannot be explained merely by examining the effect of landscape heterogeneity on animal space use (as is the norm in resource selection studies and many other species distribution models).

A central theme that runs throughout this paper is that correlative models are not sufficient for predicting space use patterns of multiple species in novel environments. This can be illustrated by a simple thought experiment. Imagine there are two populations, each of whose space use is affected by the other. One could understand the effect of population A on the space use of population B by using a correlative model, such as resource or step selection, with population B as the response variable and A as the explanatory variable. But then to predict the space use of B in a novel environment, one would need to know *a priori* the space use of A. Flipping this, one could put the distribution of A as the response variable and B as explanatory. But then predicting the space use of A requires *a priori* knowledge about B. If there is a novel environment where one does not know about the space use of either A or B then correlative models (including joint species distribution models) cannot be used for prediction. Instead a dynamic model is needed, such as an IBM or PDE. Although such IBMs and PDEs can be *parametrised* by correlative techniques (Schlägel *et al*., 2019; Potts & Schlägel, 2020), *prediction* in a multi-population situation needs techniques beyond correlation. Our purpose here is to make inroads into building these techniques.

Overall, our study aims to provide both insights into the effect of stigmergent interactions between populations on the spatio-temporal distribution of mobile species, and provide extensible methods for studying these emergent features. This complements the burgeoning statistical field of joint species distribution modelling, which gives tools for inferring the effect of one (or more) species on the distribution of another (Ovaskainen & Abrego, 2020), whilst also enhancing this field by demonstrating the importance of considering the nonlinear feedbacks between the movement processes of constituent populations for understanding spatial distributions.

## 2 Methods

### 2.1 The model

Our model of animal movement and stigmergent interactions is based on a nearest-neighbour lattice random walk formalism. We work on an *L* × *L* square lattice, Λ. We choose periodic boundary conditions for simplicity of presentation, although other forms are possible. We assume that there are *N* populations and that, for each index *i* = 1, …,*N*, population *i* consists of *M*_*i*_ individuals. Individuals leave marks at each lattice site they visit, and those marks decay geometrically over time. For simplicity, one can think of these marks as scent, such as faeces or urine, but they could correspond to any form by which animals may leave a trace of their presence on the environment. The movement of each individual is biased by the presence of marks from both their own population and others. For each population, this bias could be either attractive or repulsive, depending on whether it is beneficial or detrimental for individuals of one population to be in the presence of another population. Since animals look at their surroundings at a distance to make movement decisions, our model allows for individuals to respond to the local average density of nearby marks.

Mathematically this situation can be described by writing down the probability *f* (**x**, *t* + *τ* |**x**′, *t*) of moving from lattice site **x′** to **x** in a timestep of length *τ*. This function *f* is known as a *movement kernel*. To construct our movement kernel, we use a generalised linear model to describe the attraction to, or repulsion from, the local average density of nearby marks. A second equation is then required to describe how marks are averaged over space. Finally, the deposition and decay of marks over time is given by a third equation. We now give precise functional forms of these three equations in turn.

Letting *l* be the lattice spacing and *m*_*i*_(**x**, *t*) be the density of marks from population *i* at location **x** at time *t*, the movement kernel is given by

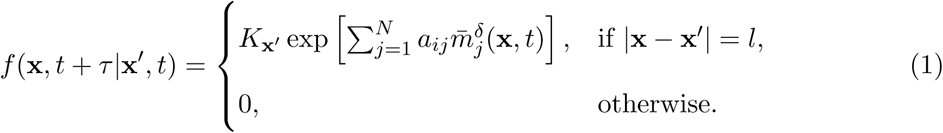

Here *K*_**x′**_ =∑_**x**_ *f* (**x**, *t* + *τ* |**x**′, *t*) is a normalising constant ensuring that *f* (**x**, *t* + *τ* |**x**′, *t*) is a well-defined probability distribution; if *a*_*ij*_ > 0 (resp. *a*_*ij*_ < 0) then |*a*_*ij*_| is the strength of population *i*’s attraction to (resp. repulsion from) population *j*; and 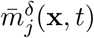 represents the average mark density over a radius of *δ*. Note that Equation (1) fits into the broad category of functions that can be parametrised by integrated step selection analysis (Avgar *et al*., 2016).

The equation for average mark density is

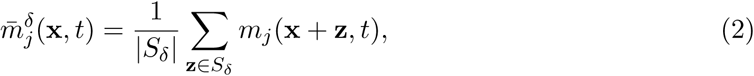

where *S*_*δ*_ = {**z** ∈ Λ : |**z**| < *δ*} is the set of lattice sites that are within a distance of *δ* from 0 and |*S*_*δ*_| is the number of lattice sites in *S*_*δ*_. Note that Equation (2) requires us to use periodic boundary conditions, so that there are always the same number of lattice sites within a distance of *δ* from any point in Λ. However, if we were to use hard boundaries, e.g. for modelling movement near a coastline, we would have to take the average in Equation (2) over the set {**x** + **z** ∈ Λ|**z** ∈ *S*_*δ*_}.

The equation defining the change in marks over time, which are deposited by individuals and then decay geometrically, is

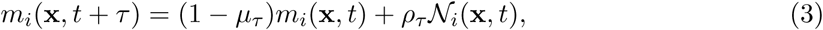

where 𝒩_*i*_(**x**, *t*) is the number of individuals at location **x** in population *i* at time *t, μ*_*τ*_ is the amount by which marks decay in a time step of length *τ*, and *ρ*_*τ*_ is the amount of marking made by a single animal in a single time step.

Equations (1-3) are not the only available functional forms to describe our stigmergent process. However, the specific form for Equation (1) is advantageous because it arrives in the form of a step selection function (Fortin *et al*., 2005; Avgar *et al*., 2016). It thus has the potential to be parametrised by the methods of Schlägel *et al*. (2019), which deals with step selection for interacting individuals (although here we focus on analysing the emergent features of the model in Equation (3) rather than the question of fitting this model to data.) Equation (2) assumes that marks are averaged over a fixed disc around the individual and was chosen for simplicity, but other options, such as exponentially decaying averaging kernels, are also possible. Equation (3) was, likewise, chosen for simplicity.

One drawback is that there is, in theory, no limit on the amount of marks in one location. If it is necessary to account for such a limit, one might exchange the *ρ*_*τ*_𝒩_*i*_(**x**, *t*) term for something like *ρ*_*τ*_ (1 − 𝒩_*i*_(**x**, *t*)*/* 𝒩_max_) 𝒩_*i*_(**x**, *t*), where 𝒩_max_ is the maximum number of marks at a single location. However, we do not explore this extension in detail here; much insight can be gained without needing to incorporate this extra complexity. Alternatively, one could replace ‘amount of marks’ with ‘probability of mark presence’. Since probabilities are bounded between 0 and 1, this would lead to a similar formalism as the situation where the number of marks has a limit. Such a situation was studied in Potts & Lewis (2016) but is not considered here.

Finally, there is an analogy between marks and resource depletion that enables our modelling framework to be used in situations where animals both deplete resources and move up resource gradients. The idea is to view the total number of marks in a location, from all the populations, as the extent of depletion of a resource. In this case, each population would avoid ‘marks’ left by either population, as animals will tend to avoid depleted resources. We do not explicitly examine this situation here, but it is a possibility for future investigations and expands the potential applicability of our work.

### 2.2 Methods for analysing simulation output

We analyse the individual-based model (IBM) from Equations (1-3) using stochastic simulations. Example simulation runs reveal a range of patterns (Fig. 2). Here, we detail methods for characterising these via three broad questions: (I) Is the distribution of animal locations heterogeneous or homogeneous? (II) If heterogeneous, do the patterns stabilise over time, so that populations keep broadly to fixed areas of space, or do they undergo persistent fluctuations? (III) For any two populations, are they segregated from one another or aggregated in the same small area? The stochastic nature of the IBM means that there will always be some amount of heterogeneity and persistent fluctuations due to noise. Our methods thus need to distinguish between what is noise and what is an actual pattern.

**Fig. 2.**
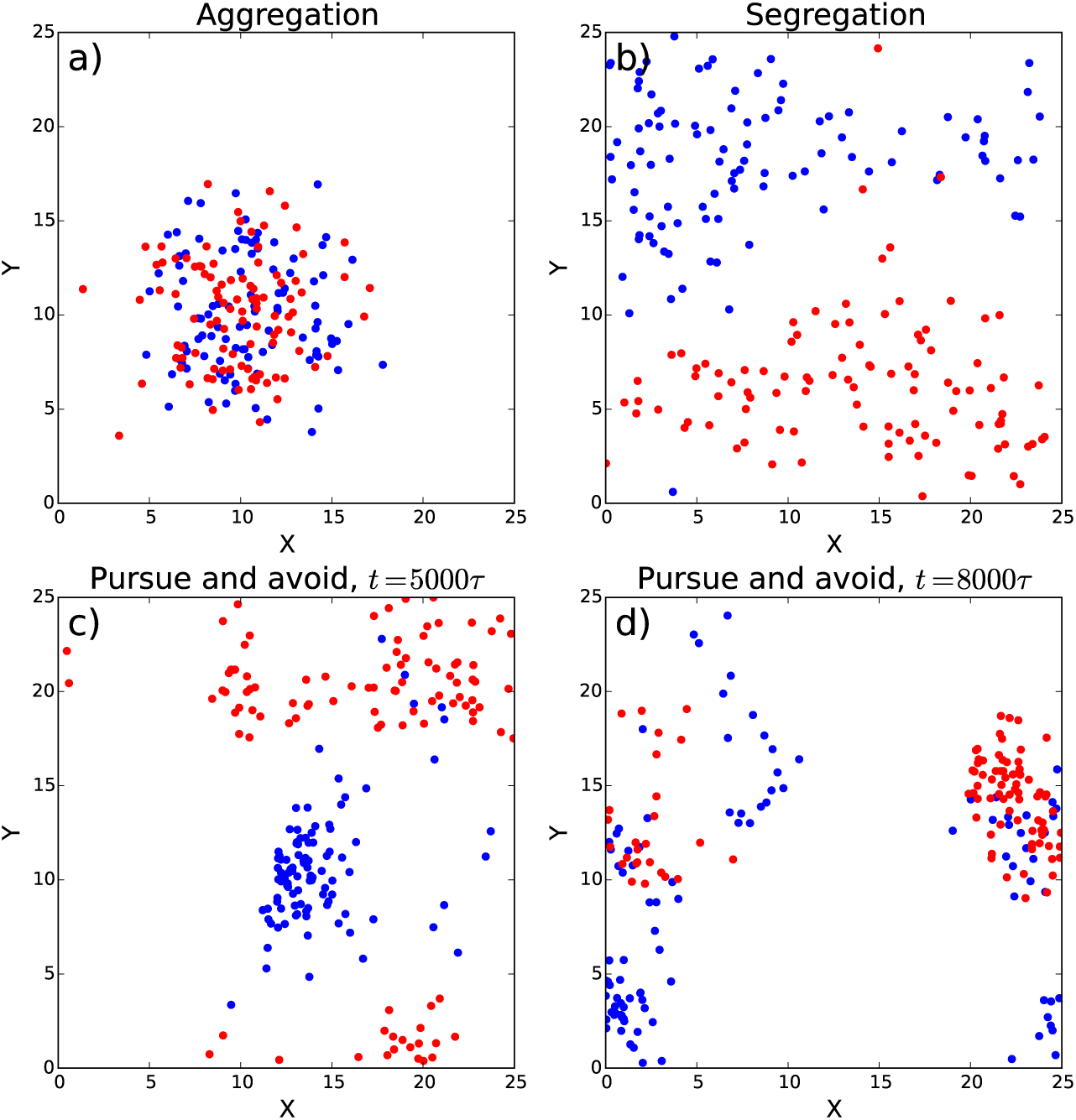
Example snapshots of simulation output. In all panels, two populations of 100 individuals each were simulated on a 25×25 lattice, with initial locations distributed uniformly at random on the landscape. Also *μ* = 0.001 and *ρ* = 0.01 for all panels (Equation 3). Panel (a) shows a system where two populations form a single, stable aggregation. Here, *a*_11_ = *a*_22_ = 0, *a*_12_ = *a*_21_ = 2, *δ* = 10*l* (Equation 1). In panel (b) the populations segregate into distinct parts of space. Here, *a*_11_ = *a*_22_ = 0, *a*_12_ = *a*_21_ = −2, and *δ* = 5*l*. In both Panels (a) and (b) the snapshot is taken at time *t* = 5000*τ*. Panels (c) and (d) show a situation where one population (blue) chases other (red) around the landscape in perpetuity, with snapshots at two different times. Here, *a*_11_ = *a*_22_ = 1, *a*_12_ = 10, *a*_21_ = −10, and *δ* = 10*l*.

To answer question (I), we examine the local population density, *l*_*i,d*_(**x**, *t*), averaged around a disc of radius *d*, at each lattice site *x* and time *t*, for each population *i*. At each point in time, we compute the *amplitude* of the pattern as *A*_*i,d*_(*t*) = max_**x**_[*l*_*i,d*_(**x**, *t*)] − min_**x**_[*l*_*i,d*_(**x**, *t*)], the maximum local population density across space minus the minimum. We want to find out whether the amplitude ever becomes higher than would be expected from individuals moving as independent random walkers (i.e. when *a*_*ij*_ = 0 for all *i, j* in Equation 1), assuming that the individuals are initially distributed uniformly at random on the lattice. For this, we calculate *A*_*i,d*_(*t*) in the case *a*_*ij*_ = 0 for all *i, j* (i.e. no mark deposition so no interactions between walkers) and take the average over a sufficiently long time period to calculate the mean to a given degree of accuracy (i.e. so that the standard deviation of the mean is below a pre-defined threshold, determined by the needs of the simulation experiment). We call this mean amplitude *A*_rw_ (for ‘random walk’). Then the extent to which the patterns are heterogenous can be determined with reference to this base-line value.

Question (II) requires that we keep track of the mean location of individuals in each population. Since individuals are moving on a lattice with periodic boundary conditions, it is necessary to take a circular mean (Berens, 2009). However, if individuals are roughly uniformly spread in either the horizontal or vertical direction then the circular mean can be very sensitive to stochastic fluctuations. We therefore introduce a *corrected circular mean* which accounts for this, and denote it by **c**_*i*_(*t*) (notice that this is a location in two dimensions, for each time, *t*). Precise details of how to calculate **c**_*i*_(*t*) are given in Supplementary Appendix A.

As with the amplitude calculations, we need to determine whether changes in **c**_*i*_(*t*) are indicative of a fluctuating pattern (like in Figs. 2c,d) or just noise around an essentially stationary population distribution (as in Figs. 2a,b). For any length *R* and time-interval, *T*, we say that a system has become (*R, T*)*-stable* at time *T*_∗_ if |**c**_*i*_(*T*_∗_ + *t*) − **c**_*i*_(*T*_∗_)| < *R* for each population *i* whenever 0 ≤ *t* ≤ *T*. For example, the systems in Figs. 2a,b are both (*l*, 1000*τ*)-stable, but the system shown in Figs. 2c,d is not. In Section 2.3 we will show how to choose values of *R* and *T*, by ensuring they are consistent with the results of mathematical analysis.

For Question (III), the extent to which a pair of populations *i, j* (*i* ≠ *j*) is aggregated or segregated at any point in time is measured using the *separation index, s*_*ij*_(*t*) = |**c**_*i*_(*t*) − **c**_*j*_(*t*)|. For systems that become (*R, T*)-stable at some time *T*_∗_, we can define the asymptotic separation index 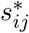 as the average of *s*_*ij*_(*t*) across *T*_∗_ < *t* < *T*_∗_ + *T*. A separation index close to 0 indicates that the populations are occupying a similar part of space. If we know, from Question (I), that both populations are displaying heterogeneous patterns then in this case we have an aggregation of both populations. Higher separation indices, coupled with the existence of heterogeneous patterns, are suggestive of segregation patterns.

The separation index is a simple metric that is quick to calculate for multiple simulation analysis. However, one could also use more sophisticated measures of range overlap, such as the Bhattacharyya’s Affinity (Fieberg & Kochanny, 2005) between kernel density estimators (Worton, 1989; Fleming *et al*., 2015). Here, though, we will keep things simple, to enable analysis of a broader range of simulation scenarios in a realistic time-frame.

### 2.3 Mathematical techniques

Techniques for analysing the output of stochastic IBMs can involve choices that might be somewhat arbitrary, for example the choices of *T*_amp_, *R*, and *T* in Section 2.2. Therefore it is valuable to ground-truth these choices by means of mathematical analysis. In particular, we do this via a PDE approximation describing the probability distribution of individuals for each population. In PDE theory, patterns can emerge when a change in parameter causes the system to switch from a situation whereby the constant steady state (corresponding to homogeneously distributed individuals) becomes unstable, leading to the distribution tending to either a non-constant steady state (heterogeneously distributed individuals), or entering a perpetually fluctuating situation. The parameter value where the switch occurs is called a bifurcation point. The nature of this bifurcation point can be ascertained by a combination of linear stability analysis (LSA) and weakly non-linear analysis. Here we focus on LSA for simplicity (which is also called Turing pattern analysis, after Turing (1952)).

To arrive at a PDE system, we take a continuous limit in both space and time, sending *l* and *τ* to 0 such that *l*^2^*/τ* tends to a finite constant, *D* > 0. This is sometimes called the diffusion limit, as *D* is a diffusion constant, but is also referred to as the parabolic limit (Hillen & Painter, 2013). If we take this limit, and also assume that infinitesimal moments beyond the second are negligible, we arrive at the following system of PDEs (see Supplementary Appendix B for details)

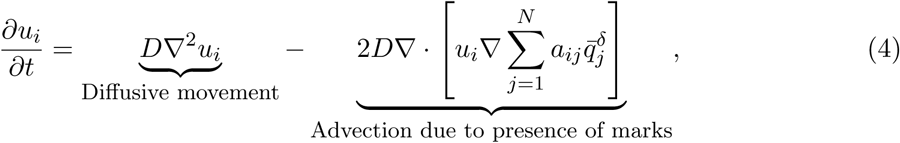

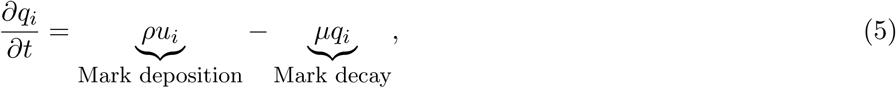

for each *i* = 1, …, *N*, where *u*_*i*_(**x**, *t*) is the location density of population *i, q*_*i*_(**x**, *t*) is the density of marks, *ρ* is the limit of 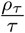 as *ρ*_*τ*_, *τ →* 0, *μ* is the limit of 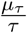 as *μ*_*τ*_, *τ →* 0, and 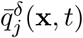 is the average of *q*(**x**, *t*) over a ball of radius *δ*. Here, we assume that animals move at the same rate, so *D* is independent of *i*. It is possible to drop this assumption, and we discuss the effect of doing this in Supplementary Appendix C. However, for simplicity of calculations we keep *D* constant in the Main Text.

It is sometimes helpful to simplify calculations by assuming that *q*_*i*_ equilibrates much faster than *u*_*i*_, so that the scent mark is in quasi-equilibrium 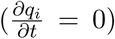, leading to the following equation for each *i* = 1, …, *N*

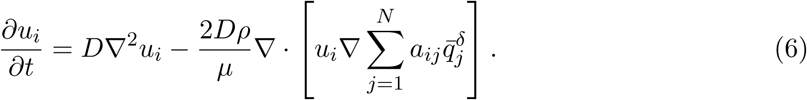

This assumption says, in effect, that the distribution of marks accurately reflects the space use distribution of the population. When terrain marking is deliberate, its usual purpose is precisely to advertise space use. Therefore this quasi-equilibrium assumption is likely to be biologically reasonable in many realistic situations.

The LSA technique enables us to use Equations (4-5) to construct the *pattern formation matrix*, ℳ (see Supplementary Appendix C for the full expression and derivation). The eigen-values of ℳ give key information about whether heterogeneous patterns will spontaneously form from small perturbations of a homogeneous system (i.e. individuals initially uniformly distributed on the landscape), and also whether these patterns begin to oscillate as they emerge. This technique dates back to Turing (1952) and is essentially an extension to PDEs of stability analysis for ordinary differential equations (May, 2019).

The emergence of heterogeneous patterns is expected whenever there is an eigenvalue whose real part is positive. Thus the sign of the eigenvalue with biggest real part (a.k.a. the dominant eigenvalue) gives an indication of the answer to Question (I) above. If the dominant eigenvalue has positive real part and a non-zero imaginary part then small perturbations of the homogeneous system will oscillate as they grow, at least at small times. Often (but not always) these oscillations will persist for all times, so give an indication of the likely answer to Question (II). We stress that this is just an indication, though, and that discrepancies may exist between the answer to (II) and whether or not the dominant eigenvalue of ℳ is real. Full analysis of when to expect non-constant stationary patterns in Equation (4-5), or when to expect perpetually changing patterns, requires more sophisticated techniques.

### 2.4 Simulation experiments

To give some insight into the sort of patterns that can emerge from our model (Equations 1-3), we perform a variety of simulations in the simple case of two populations (*N* = 2). Throughout, we assume that each population has 100 individuals (*M*_1_ = *M*_2_ = 100) and we work on a 25×25 lattice. We assume *τ* = 1 and *l* = 1 so can write *μ*_*τ*_ = *μ* and *ρ*_*τ*_ = *ρ* for ease of notation. We also assume *δ* = 5 throughout.

First, we examine the situation where populations have a symmetric response to one another, so that *a*_12_ = *a*_21_ = *a*. For simplicity, we set *a*_11_ = *a*_22_ = 0. In this case the continuum limit PDE system (Equations 4-5) has the following pattern formation matrix (derived in Supplementary Appendix C)

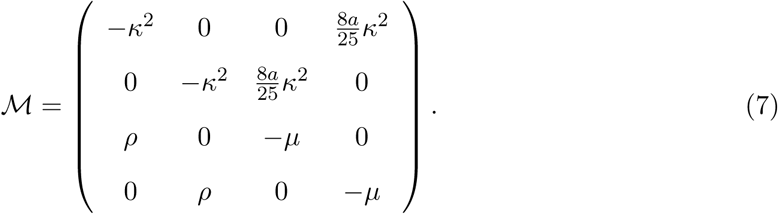

Here, *κ* is the wavenumber of the patterns that may emerge at small times, if there is an eigenvalue of ℳ with positive real part (i.e. the wavelength of these patterns would be 2*π/κ*). For our simulation experiments, we fix the scent-marking rate *ρ* = 0.01 to be a low number and vary the decay rate *μ*. We consider two different values of *a*: either *a* = 2, which corresponds to populations having a mutual attraction, or *a* = −2, corresponding to mutual avoidance. In either case, the dominant eigenvalue of ℳ is always real (Supplementary Appendix C). Furthermore, it is positive if and only if *μ* < 0.0064. In other words, this mathematical analysis predicts that the system will bifurcate at *μ* = 0.0064 from homogeneous patterns (*μ* > 0.0064) to heterogeneous patterns (*μ* < 0.0064). This means that if marks remain long enough on the landscape, they will affect movement to such an extent that the overall space use patterns change from being homogeneous (so indistinguishable from independent random walkers) to heterogeneous. This hetergogeneity will be either aggregative, if *a* = 2, analogous to the example in Figure 2a or segregative, if *a* = −2, like Figure 2b.

To test whether we see a similar change in stability in simulations, we start by simulating our system in the case *μ* = 0.009, run this until it is (*R, T*)-stable for *R* = 1 and *T* = 1000 and measure the asymptotic amplitude, 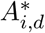 for *i* = 1, 2, by averaging *A*_*i,d*_(*t*) over the 10000 time steps after (*R, T*)-stability has been achieved. For this, we use *d* = 5. We then use the final locations of each individual as initial conditions in our next simulation run, which is identical except for choosing *μ* = 0.0069. We iterate this process, reducing *μ* by 0.0001 each time, until *μ* = 0.001. This mimics the numerical bifurcation analysis often performed when analysing PDEs (Painter & Hillen, 2011). We perform this whole iterative process for both *a* = 2 and *a* = −2, the expectation being that 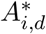 will be approximately the same as that of non-interacting individuals (*A*_rw_) until the value of *μ* crosses *μ* = 0.0064, at which point we expect 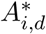 to start increasing.

To investigate whether linear stability analysis of the PDE system (Equations 4-5) reflects our method for answering Question (II), we set *a*_11_ = *a*_22_ = 1, *ρ* = 0.01, *μ* = 0.002, and sample *a*_12_ and *a*_21_ uniformly at random, 100 times each, from the interval [−5, 5]. To make calculations more transparent, we assume that the scent marks are in quasi-equilibrium, taking the adiabatic approximation in Equation (6). In this case the pattern formation matrix is

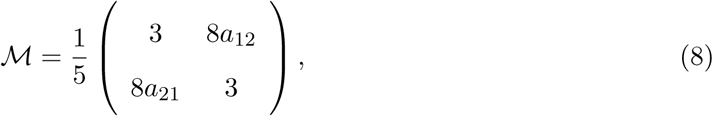

and so the dominant eigenvalue is 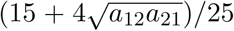. If the cross interaction terms are of identical sign (*a*_12_*a*_21_ > 0) then linear stability analysis predicts stationary patterns to emerge (at least at small times), but if they are of different sign (*a*_12_*a*_21_ < 0) then the dominant eigenvalue is not real, so patterns should oscillate as they emerge. The latter case corresponds to the type of pursuit-and-avoidance situation that we see in Fig. 2c,d. We compare these predictions to our definition of (*R, T*)-stability for a range of values of *R* and *T* to ascertain the extent to which the separation between real and non-real eigenvalues corresponds to the existence or not of (*R, T*)-stability.

### 2.5 Incorporating environmental effects

As mentioned at the end of Section 2.1, Equation (1) is in the form of a step selection function. This means that it can be readily used to incorporate the effect on movement of environmental or landscape features. Suppose that we have *n* such features, denoted by functions *Z*_1_(**x**), …, *Z*_*n*_(**x**). For each *k* = 1, …, *n*, denote by *β*_*k*_ the relative effect of *Z*_*k*_(**x**) on movement. Then, to incorporate these into the movement kernel, we modify Equation (1) as follows

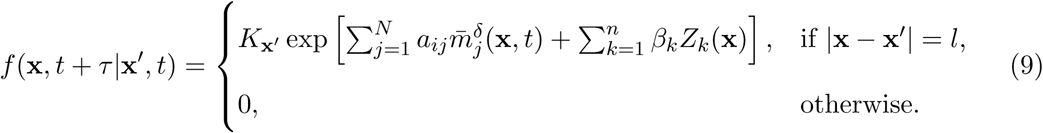

We use this to investigate the effect on space use of interactions both between populations and with the environment, by considering a simple toy scenario, but one that is based on a particular empirical situation. Specifically, we consider two populations competing for the same heterogeneously-distributed resource, *Z*_1_(**x**) (here, *n* = 1). One population is assumed to be a weaker competitor, so avoids the stronger competitor, whilst the movements of the stronger are not affected by the weaker. Both have a tendency to move towards areas of higher-density resources.

In our simulations, each population consists of 100 individuals. We examine three cases. The first is where the effect of the stronger competitor on the weaker is ignored (so animals are assumed to act independently, which mirrors many basic resource/step selection studies). The second incorporates the effect of the stronger on the weaker’s movements, but treats each individual within a population as independent from the others in the population. This mirrors some recent resource selection studies whereby the movement of one population is affected by the presence of another, e.g. Vanak *et al*. (2013); Latombe *et al*. (2014). The third assumes that the stronger population are highly territorial, so are split into five separate sub-groups, each of which exhibit strong intra-group attraction but inter-group repulsion. The simulated resource layer is a Gaussian random field on a 25 × 25 lattice, previously used in the context of resource selection by (Potts *et al*., 2014b). Precise details of the simulation experiments we performed are given in Supplementary Appendix D.

Whilst this situation is a deliberately general and simplified model, it is inspired by the particular situation of wolf-coyote coexistence in the Greater Yellowstone Ecosystem. Here, the stronger competitor is the wolf population, coyotes being weaker, and the resource layer is the distribution of where prey are likely to be found. The ability for coyotes to coexist with wolves in this system has been conjectured to emerge from the territorial structures of wolves, which include relatively large interstitial regions that may be havens for coyote (Arjo & Pletscher, 2000). If true, this means that the intra-pack attraction and inter-pack avoidance mechanisms are key to understanding the space use of wolves and coyotes. The three models presented here can be viewed as testing how the different assumptions about wolf-coyote and wolf-wolf interactions might interface with resource selection to affect their space use distributions.

## 3 Results

Fig. 3 shows the results of pattern formation analysis of our IBM. The place at which the amplitude grows higher than that of random non-interacting individuals is reasonably close to the bifurcation point predicted by linear stability analysis of the corresponding continuum PDE system. However, the latter occurs at a slightly lower value of *μ* than for the IBM indicating that a slightly lower decay rate of marks is necessary to overcome the stochastic effects and allow patterns to form. In other words, the stochasticity has a mild homogenising effect.

**Fig. 3.**
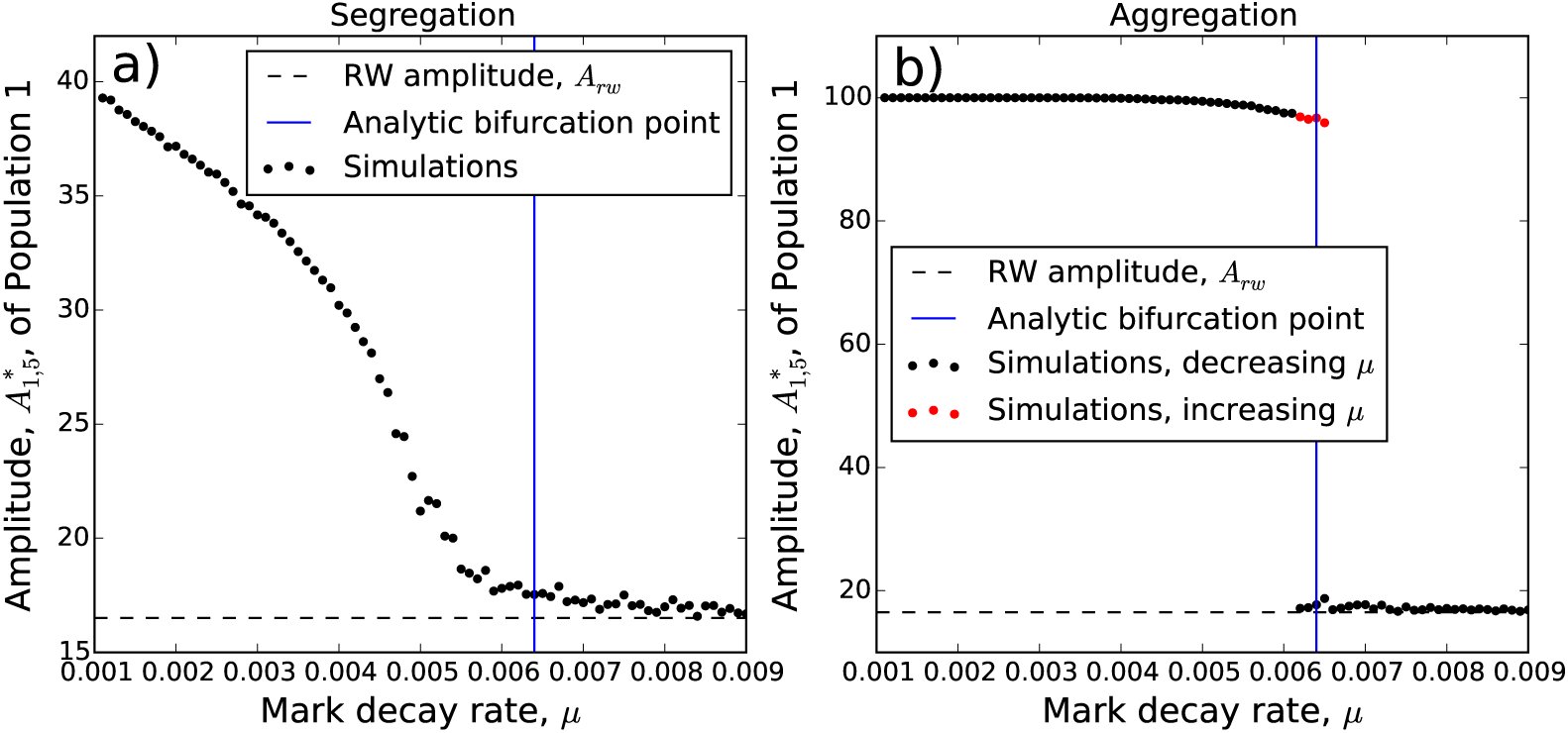
Pattern formation analysis of stochastic simulations for *N* = 2. Each panel shows, using solid dots, the amplitude, 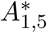, of Population 1 for different values of *μ*, where *ρ* = 0.01, *a*_12_ = *a*_21_ = *a*, and *a*_11_ = *a*_22_ = 0. Black dots represent the situation where *μ* is decreased progressively (see Section 2.4 for details) and red dots show the situation where *μ* is increased (Section 3). In Panel (a), *a* = −2 so that the populations repel one another and in Panel (b), *a* = 2 so populations are attractive. The value *A*_rw_, the amplitude in the situation where each individual is a non-interacting random walker, is given by the dashed black line. The blue line gives the bifurcation point predicted by analysis of the continuum limit PDE, Equations (4)-(5), which gives an indication of where we expect the amplitudes of the simulations to become notably larger *A*_rw_.

For negative *a* (recall *a* = *a*_12_ = *a*_21_ in Equation 1), where we tend to see segregation patterns beyond the bifurcation point, the amplitudes 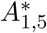, represented by black dots, appear to grow steadily as *μ* is decreased (Fig. 3a). However, for positive *a*, there is a sudden jump in the amplitude between *μ* = 0.0062 and *μ* = 0.0061 (Fig. 3b). Such jumps in bifurcation diagrams can sometimes be accompanied by a hysteresis effect, whereby if the initial conditions contain patterns then the patterns may persist even in parameter regimes where they would not emerge spontaneously. To test this, we performed the same IBM pattern formation analysis as before, but this time starting with *μ* = 0.0004 and increasing *μ* by 0.0001 each iteration (rather than decreasing as before). The red dots in Fig. 3b show that there is indeed hysteresis in the IBM system, whereby the system is bistable for 0.006 ≲ *μ* ≲ 0.0065, a phenomenon that has also been observed in single population aggregation models with a slightly different class of differential equation models (Potts & Painter, 2021). This means that if a population is already aggregated then *μ* would need to drop below about 0.006 for the aggregation to collapse. Yet if a population is not already aggregated, *μ* would have to increase above 0.0065 for aggregations to form.

Fig. 4 shows that (*R, T*)-stability corresponds well to the predictions of pattern formation analysis in the case where *R* = *l* and *T* = 5000*τ*. These were the best values of *R* and *T* we found from the ones tested, inasmuch as the results corresponded to the pattern formation analysis in the highest proportion of cases, N% (Table 1). Notice too that the mutually-avoiding populations (with *a*_12_, *a*_21_ < 0) tend to have much higher separation indices, 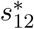, than the mutually attracting populations, as one would expect.

**Table 1.**
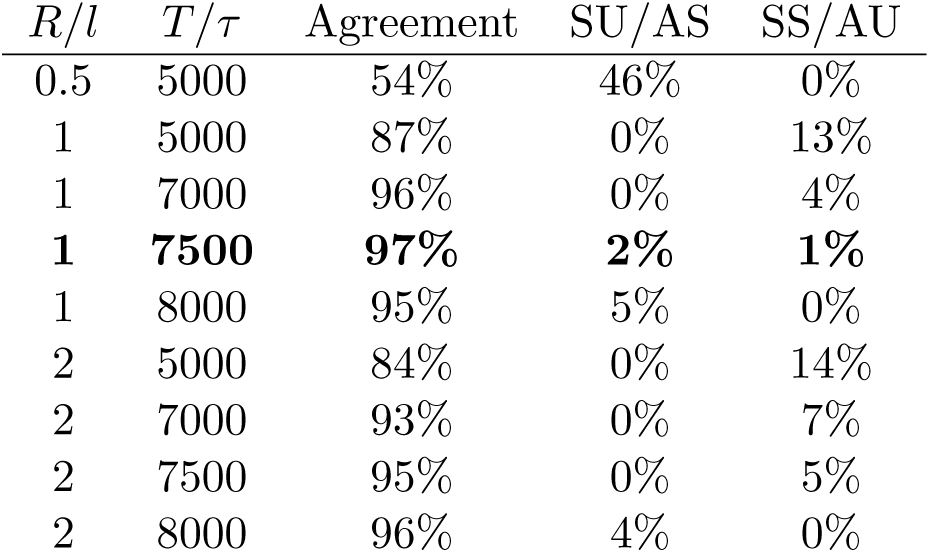
Extent to which analytic predictions agree with our simulation analysis for different choices of *R* and *T*. The third column gives the percentage of the simulations from Fig. 4 for which the analytic prediction for stability agrees with that measured from stochastic simulations using our method. The fourth (resp. fifth) gives the percentage for which the stochastic simulations were deemed unstable (resp. stable), for the given values of *R* and *T*, but the analytic prediction is stable (resp. unstable), denoted as SU/AS (rep. SS/AU).

| $R/l$ | $T/\tau$ | Agreement | SU/AS | SS/AU |
| --- | --- | --- | --- | --- |
| 0.5 | 5000 | 54% | 46% | 0% |
| 1 | 5000 | 87% | 0% | 13% |
| 1 | 7000 | 96% | 0% | 4% |
| **1** | **7500** | **97%** | **2%** | **1%** |
| 1 | 8000 | 95% | 5% | 0% |
| 2 | 5000 | 84% | 0% | 14% |
| 2 | 7000 | 93% | 0% | 7% |
| 2 | 7500 | 95% | 0% | 5% |
| 2 | 8000 | 96% | 4% | 0% |

**Fig. 4.**
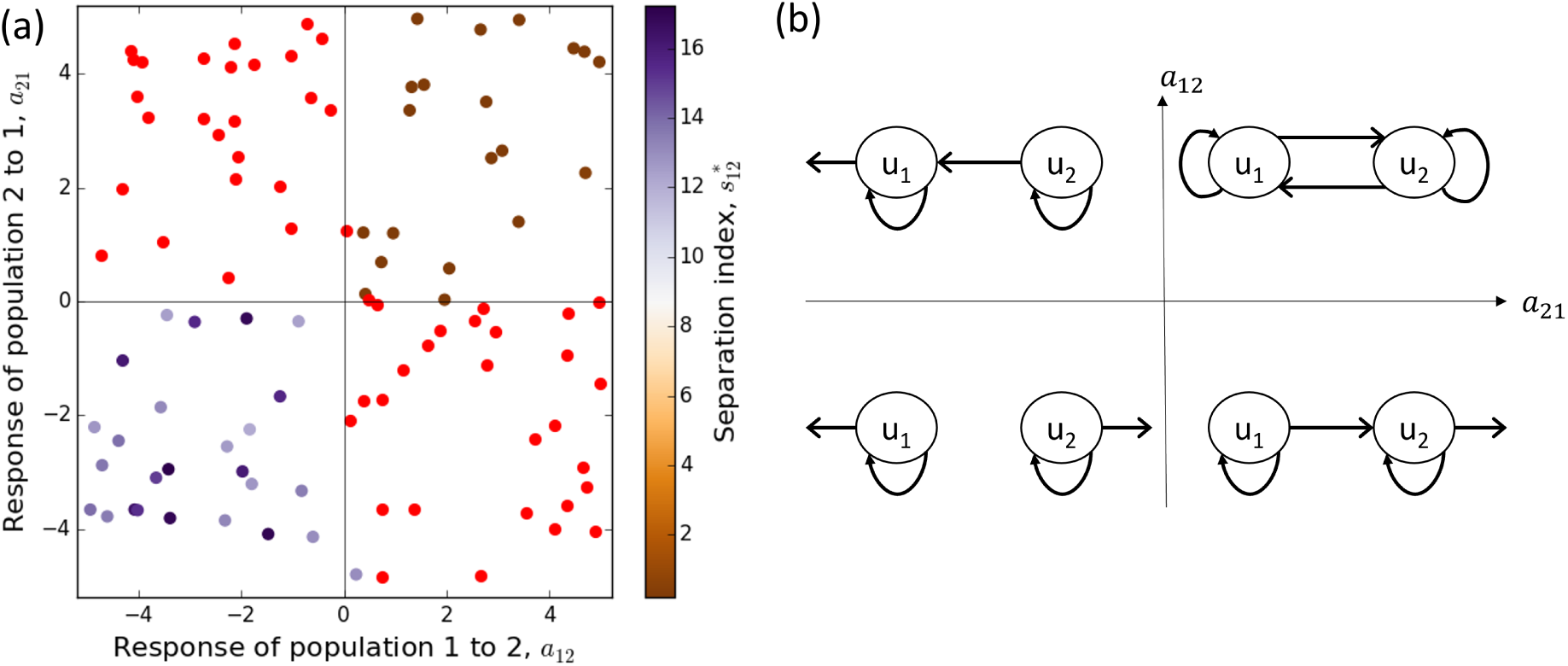
Stability of emergent patterns. In Panel (a), each dot represents a simulation run of the IBM in Equations (1)-(3) where *a*_11_ = *a*_22_ = 1, *ρ* = 0.01, *μ* = 0.002 and the values of *a*_12_ and *a*_21_ are given by the horizontal and vertical axes respectively. Red dots denote simulation runs that were not (*R, T*)-stable (for *R/l* = 1, *T/τ* = 7500), whereas those on the purple-to-brown spectrum were (*R, T*)-stable. This colour spectrum corresponds to the separation index, from aggregative to segregative. Linear pattern formation analysis of the PDEs in Equations (4)-(5) predicts stationary (resp. non-stationary) patterns to emerge in the top-right and bottom-left (resp. top-left and bottom-right) quadrants, which corresponds well with the dot colours. Notice that the top-right (resp. bottom-left) quadrant corresponds to mutual attraction (resp. avoidance) and, likewise, the dot colours indicate aggregation (resp. segregation) patterns. Panel (b) gives a schematic of the between-population movement responses corresponding to the four quadrants in panel (a). An arrow from *u*_*i*_ to *u*_*j*_ represents attraction of population *i* towards population *j*. An arrow pointing out of *u*_*i*_ away from *u*_*j*_ represents *u*_*i*_ avoiding *u*_*j*_.

Fig. 5 shows the results of our three simulation experiments on a heterogeneous resource landscape. When we assume that there are no inter-population interactions then the resulting model predicts space-use patterns whereby both populations have very similar space use distributions (Fig. 5a). When we account for the avoidance of the weaker population by the stronger then the model predicts that the stronger population will live where the resources are better, driving the weaker to resource-poor areas (Fig. 5b). This, of course, may ultimately lead to the weaker population being unable to survive. However, if the stronger population is strongly territorial, it can subdivide into separate groups, leaving interstitial regions where the weaker population can survive and have access to resources that may be relatively high quality (Fig. 5c).

**Fig. 5.**
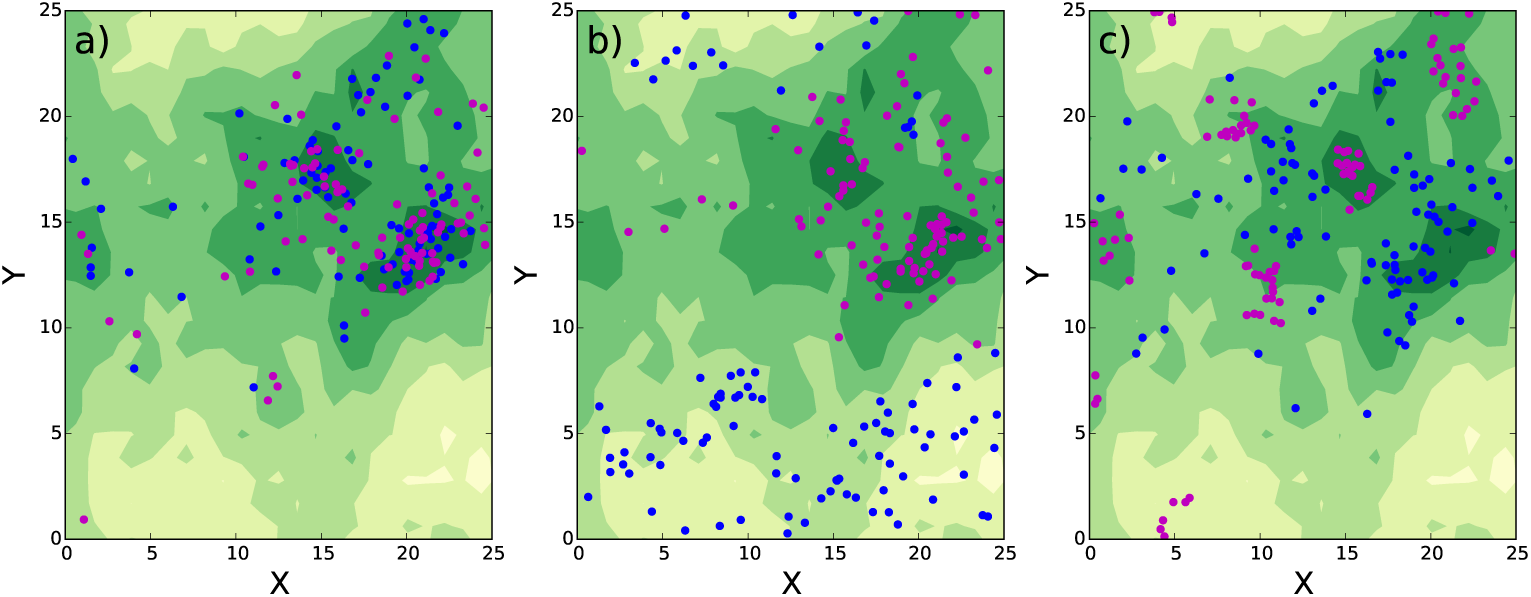
Incorporating environmental effects. This figure shows the space use distributions that emerge from three different scenarios involving two populations attracted to the same heterogeneously-distributed resource. This resource is shown in shades of yellow-green, with darker (resp. lighter) green denoting higher (resp. lower) density of resources. Magenta (resp. blue) dots denote the stronger (resp. weaker) competitor. In Panel (a) the individuals do not alter their movement in response to the presence of others, and we simply see a preference for higher quality resources. In Panel (b), as well as attraction to better resources, the weaker (blue) population has a tendency to move away from the stronger (magenta) population. In addition to this avoidance mechanism, in Panel (c) the magenta population is strongly territorial. This leads to the emergence of interstitial regions where the blue population can access resources that may be quite high quality.

## 4 Discussion

Resource selection analysis is one of the most popular techniques for understanding the distribution of species and populations. However, like many species distribution models, studies tend to focus on correlating animal locations with environmental and landscape features. Whilst some more recent studies in resource selection (Bastille-Rousseau *et al*., 2015), step selection (Vanak *et al*., 2013), and species distribution modelling (Ovaskainen & Abrego, 2020) have examined the way presence of one population may affect that of another, the first population tends to be treated as a static layer, similar to a resource layer, which then affects the presence or movement of the second population. This assumption neglects the dynamic feedbacks that can occur between two or more populations of animals.

Here, we have shown that such feedbacks can generate a variety of emergent patterns that can be quite different to those that appear when only accounting for static layers (Fig. 5). We have given a basic categorisation scheme for these patterns via simple binary questions: homogeneous or heterogeneous, stable or dynamic, segregated or aggregated. We have shown that, even with just two populations, all these patterns are possible. This categorisation, however, is likely to be only the tip of the iceberg in terms of the possible patterning properties arising from sigmergent interactions between multiple populations. Indeed, a recent study of the limiting deterministic PDE (Potts & Lewis, 2019) unveiled a rich suite of patterns through numerical simulations, including all those patterns studied here, as well as period doubling bifurcations, travelling-waves, and irregular patterns suggestive of chaos. Although such subtleties in pattern formation may be tricky to distinguish from noise in an IBM, it is valuable to be aware that they may yet be present in real systems.

Whilst a coarse-grained, individual-based approach to ecological modelling is valuable in ensuring emergent phenomena are not simply an outcome of continuum assumptions (Durrett & Levin, 1994; Getz *et al*., 2018), here our limiting PDE has been very valuable for gaining insight into our IBM. First, understanding the places where the PDE system bifurcates from between different patterning regimes has enabled us to identify interesting parameter regimes for studying our IBM (Figs. 3 and 4). Second, comparison between patterns in our IBM and the corresponding PDE has enabled us to tune the various otherwise-arbitrary choices of parameters used in analysing IBMs (e.g. the choices of *R* and *T* determined by Table 1). Whilst there is a tradition of ecological studies where limiting PDEs have helped decode the complexity inherent in IBMs (Durrett & Levin, 1994; Sherratt *et al*., 1997; Hosseini, 2006), this is perhaps overshadowed by the recent prevalence of IBM-only studies in ecology (Grimm, 1999; Grimm & Railsback, 2013; DeAngelis, 2018). We hope our use of PDEs here helps encourage further studies to implement PDEs as a tool for understanding IBMs.

Here, we have explored pattern formation analysis of PDEs using perhaps the simplest tool, that of linear analysis. However, there are plenty of other tools, with varying conceptual and mathematical complexity, that may provide insight. For example, in Fig. 3a, we see that patterns emerge smoothly as one decreases *μ* past the bifurcation point, which is suggestive of a super-critical bifurcation. However, in Fig. 3b, there is a sudden jump, together with a hysteresis (bistable) region, something usually accompanied by a sub-critical bifurcation. Techniques such as weakly non-linear analysis (Eftimie *et al*., 2009) and Crandall-Rabinowitz abstract bifurcation theory (Buttenschön & Buttenschön, 2021) are able to distinguish rigorously between these two cases. These are, however, much more conceptually and technically demanding than linear analysis, and will require a significant, separate work.

Even without advanced techniques for studying PDEs, though, we have shown how researchers can gain insight through stochastic IBM experiments. To do this, we have developed tools that mimic those used for understanding PDEs, but tailored for use with stochastic IBMs. A principal technical obstacle was to seperate-out random noise from actual spatial patterns, be they stationary or fluctuating. The fact that our techniques agreed well with the analogous PDE analysis provides a validation and ground-truthing of the methods, suggesting they are capturing the key features of patterning with good accuracy.

Furthermore, even in the relatively simple example situations studied here, our IBM analysis revealed some interesting theoretical insights. It appears that segregation patterns emerge in a continuous fashion as a parameter value moves past the bifurcation point (Figure 3a). However, when aggregations emerge, they appear suddenly (Figure 3b), with a small change in parameter value causing a sudden jump from homogeneous patterns to clearly-defined aggregations. Moreover, this is accompanied by a hysteresis effect, meaning that identical underlying processes can give rise to either aggregation or homogeneity, depending on the history of the system.

As well as using our methods to analyse IBMs, it is conceivable that the same methods may be valuable for analysing pattern formation in empirical data. One would, admittedly, need some rather high quality data: large quantities of co-tagged animals for sufficiently long time periods to observe changes in space use patterns. However, in the present ‘golden age’ of animal movement data (Wilmers *et al*., 2015), with ongoing rapid increases in the magnitude and quality of datasets (Williams *et al*., 2020), it is good idea to ensure the methodological and theoretical tools exist to deal with such data as it emerges. We have not focused on data analysis here, but we encourage researchers to test this idea in future studies if they have such data.

On the more ecological side, we have shown how accounting for feedbacks between the movement mechanisms of constituent populations may help explain the emergence of interstitial regions in territorial animals that could provide safe-havens for weaker competitors. Such patterns have been observed in coexistent wolf and coyote populations in the Greater Yellowstone Ecosystem. There, these interstitial regions have also been observed as refuges for deer (Lewis & Murray, 1993). Although we did not consider the mobility of prey resources in our simple example, one could add extra complexity by considering the attempts of mobile prey to avoid predators, and observe how this affects the spatial patterns. However, for the purposes of our simple illustration, this level of modelling complexity was not required.

An important thing to note is that emergent patterns from interacting populations cannot be revealed by correlative models alone. To take the example from Figure 5, if one knew the distribution of the stronger population, one could perform resource selection analysis with this distribution and the resource layer as the two explanatory variables to understand how these drive space use of the weaker population. However, to use this in a novel environment to predict space use of the weaker population, one would need to know *a priori* the distribution of the stronger. If one wants to predict space use of both populations at the same time, in situations where there is no *a priori* knowledge of either population, resource selection functions are not enough. Instead, one could perform step selection analysis for both interacting population, for example using the techniques of Schlägel *et al*. (2019), then feed the output of this into a movement kernel in the form of Equation (1), for example using the techniques of Potts & Schlägel (2020). This would lead to precisely the sort of IBM studied here, which enables analysis of predicted space use patterns in novel environments.

In general, our approach is valuable for predicting the distribution of populations whenever the locations of two or more populations affect the movements of each other (Schlägel *et al*., 2020). This has been observed in a variety of situations. We have already mentioned competition between carnivores, and indeed the movements of coexistent carnivore populations in response to the presence of others has been measured in various studies (Vanak *et al*., 2013; Swanson *et al*., 2016). Also the effect of predator movement on prey locations (sometimes called prey-taxis), and vice versa (the ‘landscape of fear’), has been documented in a variety of scenarios (Kareiva & Odell, 1987; Laundré *et al*., 2010; Latombe *et al*., 2014; Gallagher *et al*., 2017). Despite this, the predominant species distributions models used in ecology tend to not to account for the underlying between-population movement processes and the emergent features that they engender, even in cases where they model species jointly (Ovaskainen & Abrego, 2020). Explicit modelling of the underlying movement mechanisms, as we have done here, would help plug this gap and lead to more accurate description and forecasting of species distributions.

## Acknowledgements

JRP and VG acknowledge support of Engineering and Physical Sciences Research Council (EPSRC) grant EP/V002988/1. MAL acknowledges support from the NSERC Discovery and Canada Research Chair programs. The authors thank Jason Matthiopoulos, together with anonymous reviewers and editors, for comments that have helped greatly improve the manuscript.

## Author contributions

JRP and MAL conceived and designed the research. JRP and VG performed the research. JRP led the writing of the manuscript. All authors contributed critically to the drafts and gave final approval for publication.

