## Supplementary Appendix for "Beyond resource selection: emergent spatio-temporal distributions from animal movements and stigmergent interactions"

#### Supplementary Appendix A

In this section, we explain how to calculate the *corrected circular mean*, $\mathbf{c}_i(t)$, used in the Main Text. First, to find the mean location of a set of individuals $X_t = (\mathbf{x}_1, ..., \mathbf{x}_M) = ((x_1, y_1), ..., (x_M, y_M))$ at time $t$ in an $L \times L$ box with periodic boundary conditions, we use the *circular mean*, defined as follows

$$\alpha_x(X_t) = \frac{L}{2\pi}\text{atan2}\left[\sum_{m=1}^{M}\sin\left(\frac{2\pi x_m}{L} - \pi\right), \sum_{m=1}^{M}\cos\left(\frac{2\pi x_m}{L} - \pi\right)\right],$$
$$\alpha_y(X_t) = \frac{L}{2\pi}\text{atan2}\left[\sum_{m=1}^{M}\sin\left(\frac{2\pi y_m}{L} - \pi\right), \sum_{m=1}^{M}\cos\left(\frac{2\pi y_m}{L} - \pi\right)\right],$$
$$\alpha(X_t) = (\alpha_x(X_t), \alpha_y(X_t)). \qquad (1)$$

A problem with the above definition is that if the points in $X_t$ are sampled uniformly at random from $[0, L] \times [0, L]$, then $\alpha(X_t)$ can take any value in $[0, L] \times [0, L]$ with uniform probability density. This is because the sum (mod $L$) of $M$ uniform random variables on $[0, L]$ is itself a random variable uniformly distributed on $[0, L]$. Consequently, if we use $\alpha(X_t)$ to track the mean location of individuals, $\alpha(X_t)$ will fluctuate wildly whenever the distribution of individual locations is close to uniform. This does not reflect what we are trying to measure, which is the movement of the centre of mass of the individuals.

To correct for this, we need a way of setting the mean to $(L/2, L/2)$ whenever the points in $X_t$ are classified as sufficiently close to a uniform random distribution. It turns out that, if the points in $X_t$ are approximately uniformly distributed, the length of the vector

$$\mathbf{v}(X_t) = \left[\frac{1}{M}\sum_{m=1}^{M}\sin\left(\frac{2\pi x_m}{L}-\pi\right), \frac{1}{M}\sum_{m=1}^{M}\cos\left(\frac{2\pi x_m}{L}-\pi\right)\right] \quad (2)$$

is small, becoming arbitrarily small as $M$ increases. Therefore we can detect an approximately uniform distribution by looking for situations where $|\mathbf{v}(X_t)|$ is below some threshold, i.e.

$$|\mathbf{v}(X_t)|^2 = \left[\frac{1}{M}\sum_{m=1}^{M}\sin\left(\frac{2\pi x_m}{L}-\pi\right)\right]^2 + \left[\frac{1}{M}\sum_{m=1}^{M}\cos\left(\frac{2\pi x_m}{L}-\pi\right)\right]^2 < \epsilon^2, \quad (3)$$

for some $\epsilon$. We found that, where $M = 100$, a value of $\epsilon = 0.3$ worked well for classifying distributions as uniform or otherwise. Lower values of $\epsilon$ tended to lead to situations where samples from uniform distributions were misclassified as non-uniform. We denote the resulting estimate of the centre of mass of $X_t$ as $C_\epsilon(X_t) \in [0, L] \times [0, L]$, so that $C_\epsilon(X_t) = \alpha(X_t)$ if $|\mathbf{v}(X_t)| \geq \epsilon$ and $C_\epsilon(X_t) = (L/2, L/2)$ otherwise.

Finally, to construct the corrected circular mean, we need to "unwrap" the periodic boundary conditions. The idea is that we assume that $\mathbf{c}_i(0) \in [0, L] \times [0, L]$ but if there is a point in time where $C_\epsilon(X_t) = (C_{\epsilon,x}(X_t), C_{\epsilon,y}(X_t))$ crosses the boundary, we move into the next square along in $\mathbb{R}^2$. In other words, we set $\mathbf{c}_i(t) = C_\epsilon(X_t)$ until we reach a time $t_1$ where one of the following occurs for the first time:

1. $C_{\epsilon,x}(X_{t_1}) < 1$ (resp. $C_{\epsilon,y}(X_{t_1}) < 1$) but $C_{\epsilon,x}(X_{t_1+\tau}) > L-1$ (resp. $C_{\epsilon,y}(X_{t_1+\tau}) > L-1$),
2. $C_{\epsilon,x}(X_{t_1}) > L-1$ (resp. $C_{\epsilon,y}(X_{t_1}) > L-1$) but $C_{\epsilon,x}(X_{t_1+\tau}) < 1$ (resp. $C_{\epsilon,y}(X_{t_1+\tau}) < 1$).

Denoting $\mathbf{c}_i(t) = (c_{i,x}(t), c_{i,y}(t))$, in the first case we set $c_{i,x}(t) = C_{\epsilon,x}(X_t) - L$ (resp. $c_{i,y}(t) = C_{\epsilon,y}(X_t) - L$) and in the second case, we set $c_{i,x}(t) = C_{\epsilon,x}(X_t) + L$ (resp. $c_{i,y}(t) = C_{\epsilon,y}(X_t) + L$) for any $t \geq t_1 + \tau$ until such a time as $C_\epsilon(X_t)$ crosses the boundary again, given by either situation (1) or (2) listed above. Then, each time a boundary is crossed, we repeat this process of adding or subtracting $L$ to $c_{i,x}(t)$ or $c_{i,y}(t)$, depending on which boundary is crossed.

### Supplementary Appendix B

In this section, we explain how to derive the PDE in Equations (4-5) of the Main Text, from the IBM given in Equations (1-3) of the Main Text. Recall the following equations for the IBM (see the Main Text for definitions of the various terms)

$$f(\mathbf{x}, t+\tau|\mathbf{x}', t) = \begin{cases} K_{\mathbf{x}'} \exp\left[\sum_{j=1}^{N} a_{ij}\bar{m}_j^{\delta}(\mathbf{x}, t)\right], & \text{if } |\mathbf{x}-\mathbf{x}'| = l, \\ 0, & \text{otherwise.} \end{cases} \tag{4}$$

$$\bar{m}_j^{\delta}(\mathbf{x}, t) = \frac{1}{|S_\delta|} \sum_{\mathbf{z}\in S_\delta} m_j(\mathbf{x}+\mathbf{z}, t), \tag{5}$$

$$m_i(\mathbf{x}, t+\tau) = (1-\mu_\tau)m_i(\mathbf{x}, t) + \rho_\tau \mathcal{N}_i(\mathbf{x}, t). \tag{6}$$

Beginning with Equation (6), let $Q_i(\mathbf{x}, t)$ be the expected value of $m_i(\mathbf{x}, t)$ and $P_{i,k}(\mathbf{x}, t)$ be the probability of individual $k$ from population $i$ being at location $\mathbf{x}$ at time $t$. Then taking expectations of Equation (6) gives

$$Q_i(\mathbf{x}, t+\tau) = (1-\mu_\tau)Q_i(\mathbf{x}, t) + \rho_\tau \sum_{k=1}^{M_i} P_{i,k}(\mathbf{x}, t). \tag{7}$$

Subtracting $Q_i(\mathbf{x}, t)$ from both sides and dividing by $\tau$ gives

$$\frac{Q_i(\mathbf{x}, t+\tau) - Q_i(\mathbf{x}, t)}{\tau} = \frac{\rho_\tau}{\tau} \sum_{k=1}^{M_i} P_{i,k}(\mathbf{x}, t) - \frac{\mu_\tau}{\tau} Q_i(\mathbf{x}, t). \tag{8}$$

Then we take the limit as $l, \tau, \rho_\tau, \mu_\tau \to 0$ such that $(\rho_\tau/\tau) \to \rho$ and $(\mu_\tau/\tau) \to \mu$. Writing $q_i(\mathbf{x}, t)$ and $u_{i,k}(\mathbf{x}, t)$ for the density functions corresponding to $Q_i$ and $P_{i,k}$ respectively in this limit, we arrive at

$$\frac{\partial q_i}{\partial t} = \rho u_i - \mu q_i, \tag{9}$$

which is Equation (5) from the Main Text.

To derive the PDE governing the evolution of $u_i(\mathbf{x}, t)$ (Equation 4 from the Main Text), we first re-write Equation (4) as follows

$$f(\mathbf{x}, t+\tau|\mathbf{x}', t) = K_{\mathbf{x}'}\phi_\tau(\mathbf{x}-\mathbf{x}')\exp\left[\sum_{j=1}^{N} a_{ij}\bar{m}_j^\delta(\mathbf{x}, t)\right], \tag{10}$$

$$\phi_\tau(\mathbf{z}) = \frac{1}{4}\sum_{\mathbf{z}'\in S}\delta(\mathbf{z}-\mathbf{z}'), \tag{11}$$

where $\delta$ denotes the Dirac delta function and $S\{(-l,0),(l,0),(0,-l),(0,l)\}$. Notice that $\bar{m}_j^\delta(\mathbf{x}, t) \approx \bar{q}_i(\mathbf{x}, t)$, the latter being the expectation of the former, so we have

$$f(\mathbf{x}, t+\tau|\mathbf{x}', t) \approx K_{\mathbf{x}'}\phi_\tau(\mathbf{x}-\mathbf{x}')\exp\left[\sum_{j=1}^{N} a_{ij}\bar{q}_j(\mathbf{x}, t)\right]. \tag{12}$$

We then use a result from Potts & Schlägel (2020, Equations 3-4), which uses an old moment closure result from Patlak (1953) to show that $u_i(\mathbf{x}, t)$ is approximately governed by the following PDE

$$\frac{\partial u_i}{\partial t} = D_{l,\tau}\nabla^2 u_i - 2D_{l,\tau}\nabla\cdot\left[u_i\nabla\sum_{j=1}^{N} a_{ij}\bar{q}_j\right], \tag{13}$$

where

$$D_{l,\tau} = \frac{1}{4\tau}\int_{\mathbb{R}^2}|\mathbf{x}|^2\phi_\tau(|\mathbf{x}|)\mathrm{d}\mathbf{x} = \frac{l^2}{4\tau}. \tag{14}$$

Letting $D$ be the limit of $D_{l,\tau}$ as $l, \tau \to 0$, Equation (13) becomes

$$\frac{\partial u_i}{\partial t} = D\nabla^2 u_i - 2D\nabla\cdot\left[u_i\nabla\sum_{j=1}^{N} a_{ij}\bar{q}_j\right], \tag{15}$$

which is Equation (4) from the Main Text.

### Supplementary Appendix C

Here we explain how the various pattern formation matrices from the Main Text were calculated. The PDE system from Equations (4-5) in the Main Text is as follows

$$\frac{\partial u_i}{\partial t} = \underbrace{D\nabla^2 u_i}_{\text{Diffusive movement}} - \underbrace{2D\nabla \cdot \left[ u_i \nabla \sum_{j=1}^{N} a_{ij} \bar{q}_j^{\delta} \right]}_{\text{Advection due to presence of marks}}, \tag{16}$$

$$\frac{\partial q_i}{\partial t} = \underbrace{\rho u_i}_{\text{Mark deposition}} - \underbrace{\mu q_i}_{\text{Mark decay}}, \tag{17}$$

The technique of linear pattern formation analysis is very standard within the mathematical literature. It began with Turing (1952), but can be found in many textbooks, such as Murray (2003), which is an excellent reference for the interested novice with a biological leaning. The idea is to look at solutions that are small, spatially non-constant, perturbations of the constant steady state and see if they grow or shrink over time. Growing perturbations are indicative of spontaneous pattern formation.

In our case, the constant steady state is given by

$$u_i^* = \frac{1}{L^2} \int_0^L \int_0^L u_i(x, y, t) \mathrm{d}x \mathrm{d}y, \tag{18}$$

$$q_i^* = \frac{\rho}{\mu} u_i^*, \tag{19}$$

where we have written $(x, y)$ for $\mathbf{x}$. Observe that $u_i(x, y, t) = u_i^*$ and $q_i(x, y, t) = q_i^*$ are steady states of Equations (16-17). We look for approximate solutions of the form $u_i(x, y, t) = u_i^* + \hat{u}_i(x, y, t)$ and $q_i(x, y, t) = q_i^* + \hat{q}_i(x, y, t)$ where $\hat{u}_i(x, y, t) = u_{i,0} \exp(\sigma t + \mathrm{i}\kappa_x x + \mathrm{i}\kappa_y y)$ and $\hat{q}_i(x, y, t) = q_{i,0} \exp(\sigma t + \mathrm{i}\kappa_x x + \mathrm{i}\kappa_y y)$, for arbitrarily small constants $u_{i,0}$ and $q_{i,0}$. Neglecting non-linear terms, this ansatz leads to the following eigenvector equation

$$\sigma \begin{pmatrix} \hat{u}_1 \\ \hat{u}_2 \\ \vdots \\ \hat{u}_N \\ \hat{q}_1 \\ \hat{q}_2 \\ \vdots \\ \hat{q}_N \end{pmatrix} = \begin{pmatrix} -\kappa_x^2 D & 0 & \dots & 0 & 2\kappa_x^2 D u_1^* a_{11}\mathrm{sinc}(\delta\kappa_x) & \dots & 2\kappa_x^2 D u_1^* a_{1N}\mathrm{sinc}(\delta\kappa_x) \\ \vdots & \vdots & \dots & \vdots & \vdots & \dots & \vdots \\ 0 & 0 & \dots & -\kappa_x^2 D & 2\kappa_x^2 D u_N^* a_{N1}\mathrm{sinc}(\delta\kappa_x) & \dots & 2\kappa_x^2 D u_N^* a_{NN}\mathrm{sinc}(\delta\kappa_x) \\ \rho & 0 & \dots & 0 & -\mu & \dots & 0 \\ \vdots & \vdots & \dots & \vdots & \vdots & \dots & \vdots \\ 0 & 0 & \dots & \rho & 0 & \dots & -\mu \end{pmatrix} \begin{pmatrix} \hat{u}_1 \\ \hat{u}_2 \\ \vdots \\ \hat{u}_N \\ \hat{q}_1 \\ \hat{q}_2 \\ \vdots \\ \hat{q}_N \end{pmatrix}$$

$$+ \begin{pmatrix} -\kappa_y^2 D & 0 & \dots & 0 & 2\kappa_y^2 D u_1^* a_{11}\mathrm{sinc}(\delta\kappa_y) & \dots & 2\kappa_y^2 D u_1^* a_{1N}\mathrm{sinc}(\delta\kappa_y) \\ \vdots & \vdots & \dots & \vdots & \vdots & \dots & \vdots \\ 0 & 0 & \dots & -\kappa_y^2 D & 2\kappa_y^2 D u_N^* a_{N1}\mathrm{sinc}(\delta\kappa_y) & \dots & 2\kappa_y^2 D u_N^* a_{NN}\mathrm{sinc}(\delta\kappa_y) \\ \rho & 0 & \dots & 0 & -\mu & \dots & 0 \\ \vdots & \vdots & \dots & \vdots & \vdots & \dots & \vdots \\ 0 & 0 & \dots & \rho & 0 & \dots & -\mu \end{pmatrix} \begin{pmatrix} \hat{u}_1 \\ \hat{u}_2 \\ \vdots \\ \hat{u}_N \\ \hat{q}_1 \\ \hat{q}_2 \\ \vdots \\ \hat{q}_N \end{pmatrix}, \tag{20}$$

where $\mathrm{sinc}(x)$ is the function defined by $\sin(x)/x$ for $x \neq 0$ and $\mathrm{sinc}(0) = 1$ (notice that $\mathrm{sinc}(x)$ is infinitely differentiable). Since $\mathrm{sinc}(0) = 1$ and $\max\{\mathrm{sinc}(x)\} = 1$, if an eigenvalue $\sigma$ has positive real part for some $\delta$, $\kappa_x$, and $\kappa_y$ then it will be positive for arbitrarily small $\delta$. Thus, taking the limit $\delta \to 0$ and writing $\kappa^2 = \kappa_x^2 + \kappa_y^2$, we have

$$\sigma \mathbf{w} = \mathcal{M}\mathbf{w},$$

$$\mathbf{w} = \begin{pmatrix} \hat{u}_1 \\ \hat{u}_2 \\ \vdots \\ \hat{u}_N \\ \hat{q}_1 \\ \hat{q}_2 \\ \vdots \\ \hat{q}_N \end{pmatrix}, \quad \mathcal{M} = \begin{pmatrix} -\kappa^2 D & 0 & \dots & 0 & 2\kappa^2 D u_1^* a_{11} & \dots & 2\kappa^2 D u_1^* a_{1N} \\ \vdots & \vdots & \dots & \vdots & \vdots & \dots & \vdots \\ 0 & 0 & \dots & -\kappa^2 D & 2\kappa^2 D u_N^* a_{N1} & \dots & 2\kappa^2 D u_N^* a_{NN} \\ \rho & 0 & \dots & 0 & -\mu & \dots & 0 \\ \vdots & \vdots & \dots & \vdots & \vdots & \dots & \vdots \\ 0 & 0 & \dots & \rho & 0 & \dots & -\mu \end{pmatrix}. \tag{21}$$

In the Main Text, we refer to $\mathcal{M}$ as the *pattern formation matrix* of the system (Equations 16-17). To calculate the specific pattern formation matrix given in Equation (7) of the Main Text, we first set $N = 2$, $D = 1$, $a_{11} = a_{22} = 0$, and $a_{12} = a_{21} = a$ to give

$$\mathcal{M} = \begin{pmatrix} -\kappa^2 & 0 & 0 & 2au_1^*\kappa^2 \\ 0 & -\kappa^2 & 2au_2^*\kappa^2 & 0 \\ \rho & 0 & -\mu & 0 \\ 0 & \rho & 0 & -\mu \end{pmatrix}. \tag{22}$$

Then we need to calculate $u_1^*$ and $u_2^*$, which are the population densities of populations 1 and 2 respectively. Since there are, in the simulation experiments from the main text, 100 individuals in a box of size $25 \times 25$, the population density is $u_1^* = u_2^* = 4/25$, which leads to

$$\mathcal{M} = \begin{pmatrix} -\kappa^2 & 0 & 0 & \frac{8a}{25}\kappa^2 \\ 0 & -\kappa^2 & \frac{8a}{25}\kappa^2 & 0 \\ \rho & 0 & -\mu & 0 \\ 0 & \rho & 0 & -\mu \end{pmatrix}, \tag{23}$$

which is Equation (7) from the Main Text. The eigenvalues of $\mathcal{M}$ are the values of $\sigma$ that satisfy

$$(\kappa^2 + \sigma)^2(\mu + \sigma)^2 - \left(\frac{8a\rho\kappa^2}{25}\right)^2 = 0. \tag{24}$$

The point at which we bifurcate from no patterns to patterns is the set of parameters that solve Equation (24) for $\sigma = 0$. This is where $\mu = \frac{8|a|\rho}{25}$. In the simulation experiments from the Main Text, we have set $\rho = 0.01$ and $a = \pm 2$. Therefore the bifurcation point is $\mu = 0.0064$, as claimed in the Main Text.

Note that here we have assumed the movement rates of the two populations are identical. If we drop this assumption then this means $D$ is replaced by $D_i$ in Equation (16). Following the calculations though, we arrive at the following version of Equation (22)

$$\mathcal{M} = \begin{pmatrix} -D_1\kappa^2 & 0 & 0 & \frac{8a}{25}D_1\kappa^2 \\ 0 & -D_2\kappa^2 & \frac{8a}{25}D_2\kappa^2 & 0 \\ \rho & 0 & -\mu & 0 \\ 0 & \rho & 0 & -\mu \end{pmatrix}. \tag{25}$$

Then the eigenvalues satisfy

$$(D_1D_2\kappa^2 + \sigma)^2(\mu + \sigma)^2 - \left(\frac{8aD_1D_2\rho\kappa^2}{25}\right)^2 = 0. \tag{26}$$

Then the demarcation between patterns and no patterns depends on $d$ as well as the other parameters. However, the techniques for analysing this more general situation remain the same.

### Supplementary Appendix D

Here we give details of the model used to construct Fig. 5 from the Main Text, which supplement the general description given in the Methods section of the Main Text. The Gaussian random field used for the resource layer was generated by the R function `GaussRF()` from the `RandomFields` package (Schlather *et al.*, 2016), using the exponential model with mean = 0, variance = 1, nugget = 0, and scale = 10.

The simulations consisted of 10 populations ($N = 10$), the first five of which can be thought of as groups of the stronger competitor and the second five as the weaker competitor. Each population consisted of 20 individuals ($M_i = 20$ for all $i$). The matrix $\mathcal{A} = (a_{ij})_{i,j}$ is given by

$$\mathcal{A} = \begin{pmatrix} 0 & 0 & 0 & 0 & 0 & 0 & 0 & 0 & 0 & 0 \\ 0 & 0 & 0 & 0 & 0 & 0 & 0 & 0 & 0 & 0 \\ 0 & 0 & 0 & 0 & 0 & 0 & 0 & 0 & 0 & 0 \\ 0 & 0 & 0 & 0 & 0 & 0 & 0 & 0 & 0 & 0 \\ 0 & 0 & 0 & 0 & 0 & 0 & 0 & 0 & 0 & 0 \\ -2 & -2 & -2 & -2 & -2 & 0 & 0 & 0 & 0 & 0 \\ -2 & -2 & -2 & -2 & -2 & 0 & 0 & 0 & 0 & 0 \\ -2 & -2 & -2 & -2 & -2 & 0 & 0 & 0 & 0 & 0 \\ -2 & -2 & -2 & -2 & -2 & 0 & 0 & 0 & 0 & 0 \\ -2 & -2 & -2 & -2 & -2 & 0 & 0 & 0 & 0 & 0 \end{pmatrix} \tag{27}$$

for the model leading to Panel 5b from the Main Text and

$$
\mathcal{A} = \begin{pmatrix}
2 & -2 & -2 & -2 & -2 & 0 & 0 & 0 & 0 & 0 \\
-2 & 2 & -2 & -2 & -2 & 0 & 0 & 0 & 0 & 0 \\
-2 & -2 & 2 & -2 & -2 & 0 & 0 & 0 & 0 & 0 \\
-2 & -2 & -2 & 2 & -2 & 0 & 0 & 0 & 0 & 0 \\
-2 & -2 & -2 & -2 & 2 & 0 & 0 & 0 & 0 & 0 \\
-2 & -2 & -2 & -2 & -2 & 0 & 0 & 0 & 0 & 0 \\
-2 & -2 & -2 & -2 & -2 & 0 & 0 & 0 & 0 & 0 \\
-2 & -2 & -2 & -2 & -2 & 0 & 0 & 0 & 0 & 0 \\
-2 & -2 & -2 & -2 & -2 & 0 & 0 & 0 & 0 & 0 \\
-2 & -2 & -2 & -2 & -2 & 0 & 0 & 0 & 0 & 0
\end{pmatrix} \tag{28}
$$

for the model leading to Panel 5c from the Main Text. We used $\delta = 5l$ to model the way the first five populations (the stronger competitor) responded to marks and $\delta = l$ for the second five populations (the weaker competitor).
